# Divergent intratumoral associations of EGFR, ERBB3 (HER3) and TACSTD2 (TROP2) transcripts with partial epithelial–mesenchymal transition in head and neck squamous cell carcinoma

**DOI:** 10.64898/2026.11.23.768765

**Authors:** Yuelong Chen, Peng Yan, Simiao Fan, Jiawei Li, Hongjian Yu, Lixin Cheng, Weidong Tang, Jijun Tang, Nana Jin

## Abstract

**Objectives:** To define how EGFR, ERBB3 (HER3) and TACSTD2 (TROP2) transcripts relate to partial epithelial–mesenchymal transition (pEMT) and epithelial differentiation within head and neck squamous cell carcinoma (HNSCC).

**Materials and methods:** Four public single-cell cohorts included 26 patients eligible for within-patient malignant-cell analyses. Target-excluded state scores defined upper and lower pEMT thirds; paired between-target expression contrasts were tested using sign-flip tests and Benjamini–Hochberg correction. Spatial, paired RNA–protein, bulk-tumor and synthetic-mixture analyses provided complementary evidence.

**Results:** EGFR showed a more positive pEMT association than ERBB3 and TACSTD2. Discovery contrasts were 1.58 log□ units (95% CI, 0.87–2.42; n = 9) and 1.30 (0.75–1.91; n = 10), respectively (both q = 0.006). Both contrasts replicated independently (both q = 0.047), with directional concordance in two additional cohorts. Differentiation showed the opposite relative pattern; its adjustment attenuated EGFR–TACSTD2 separation. Visium tumor regions from 10 patients supported both relative EGFR–pEMT associations (both q = 0.004). Paired regions from 22 patients characterized tumor buds as pEMT-high and less differentiated. CPTAC RNA–protein correlations were 0.862 for EGFR, 0.764 for TROP2 and 0.404 for HER3. EGFR tracked bulk pEMT and HER3/TROP2 tracked differentiation at both molecular levels. An EGFR-relative transcript-balance index correlated with bulk pEMT in 273 TCGA tumors (purity-adjusted ρ = 0.453).

**Conclusions:** Reproducible cell-state-associated transcript differences support evaluating therapeutic-target abundance together with independent state markers. These findings inform protein-focused biomarker studies of EGFR-, HER3- and TROP2-directed therapies in HNSCC.

## Introduction

The clinical benefit of cetuximab established epidermal growth factor receptor (EGFR) as a therapeutic target in head and neck squamous cell carcinoma (HNSCC)^1,2^. Antibody–drug conjugates (ADCs) have extended interest in EGFR, human epidermal growth factor receptor 3 (HER3) and trophoblast cell-surface antigen 2 (TROP2). Antitumor activity has been reported for the EGFR-directed ADC becotatug vedotin and the TROP2-directed ADC sacituzumab govitecan in previously treated recurrent or metastatic HNSCC^3,4^. EGFR–HER3 bispecific ADCs have also entered clinical evaluation in advanced solid tumors^5^. Quantitative tissue assays demonstrate broad but variable expression of these targets in HNSCC^6^. These developments make it important to establish not only how much target is present in a tumor, but also which malignant-cell populations express it. Malignant cells within HNSCC occupy diverse transcriptional states. Single-cell studies identified partial epithelial–mesenchymal transition (pEMT) and epithelial differentiation as prominent programs of intratumoral variation. The pEMT program combines retained epithelial features with expression of genes involved in extracellular matrix interactions and invasion^7^. Cells expressing this program have been localized to the leading edge of primary tumors, and pEMT markers have been associated with nodal metastasis in oral cavity squamous cell carcinoma^7,8^. Spatial transcriptomic studies have further identified invasion-related expression programs in histologically defined tumor buds^9^. These observations provide a rationale for examining therapeutic targets in relation to pEMT and epithelial differentiation, rather than treating malignant cells within a tumor as a uniform target-positive population.

Experimental evidence connects the three targets to epithelial plasticity in different ways. EGFR-associated regulatory networks contribute to local invasion in HNSCC^10^, whereas TROP2 loss promotes mesenchymal features in squamous carcinoma models and can activate neuregulin-dependent HER3 signaling in HNSCC^11,12^. These findings establish biological links, but do not determine how EGFR, ERBB3 (HER3) and TACSTD2 (TROP2) transcripts compare across malignant-cell states within individual human tumors. Broad tissue-level expression could coexist with divergent state-associated profiles. The unresolved question is therefore whether the three transcripts show shared or divergent associations with pEMT within the same tumor, and how these associations relate to epithelial differentiation. Addressing this question requires direct comparison of the targets within patients, rather than separate assessments of each target across tumors. Complementary molecular and spatial measurements can place these within-tumor comparisons in a translational context. Single-cell profiling resolves expression differences among malignant-cell states, while spatial profiling examines their counterparts in tissue regions. Paired RNA and protein measurements provide an additional perspective for evaluating whether transcriptional associations have protein-level counterparts^13^. Bulk mesenchymal signatures can also reflect stromal contributions, making cellular composition an important consideration when interpreting tissue-level target–state associations^14^. Considering these measurements together offers an opportunity to relate therapeutic-target expression to malignant-cell state, tissue organization and protein abundance.

Here, we investigated the relative associations of EGFR, ERBB3 and TACSTD2 transcripts with pEMT and epithelial differentiation in HNSCC, with primary single-cell comparisons focused on tumors classified as human papillomavirus (HPV)-negative or p16-negative. We reanalyzed public single-cell cohorts using direct within-patient comparisons and evaluated the reproducibility of target-specific state associations across cohorts. We complemented these analyses with spatial profiling of tumor regions and tumor buds, paired bulk RNA–protein measurements, and analyses of tissue-level expression and cellular composition. Our aim was to define the cell-state context of therapeutic-target heterogeneity and provide a biological foundation for biomarker studies of EGFR-, HER3- and TROP2-directed therapies.

## Methods

### Study design and data sources

We reanalyzed publicly available, de-identified data to compare the associations of *EGFR*, *ERBB3* and *TACSTD2* with partial epithelial–mesenchymal transition (pEMT) and epithelial differentiation in head and neck squamous cell carcinoma (HNSCC). Single-cell analyses used GSE181919 as the discovery cohort, GSE172577 as the replication cohort, and GSE188737 and GSE182227 as two additional cohorts. Spatial analyses used Visium data from GSE208253 and GeoMx data from GSE300414^9,15–19^. CPTAC-HNSCC provided paired RNA and proteomic measurements, and TCGA-HNSC provided bulk expression and clinical data^20–22^. Cohort characteristics and analytical roles are summarized in Table 1.

**Table 1.** Overview of datasets and their analytical roles.

| Dataset | Assay | Analysis population | Anatomical sites | HPV ascertainment | Analytical role |
|---|---|---|---|---|---|
| GSE181919 | scRNA-seq | 10 patients | Oral cavity; oropharynx; hypopharynx | Study-reported HPV-negative | Cellular atlas and discovery of within-patient target–state contrasts |
| GSE172577 | scRNA-seq | 6 patients | Oral cavity | HPV-negative according to the secondary atlas | Replication of within-patient target–state contrasts |
| GSE188737 | scRNA-seq | 6 patients | Oral cavity | Study-reported HPV-negative; discordant p16 status in one patient | Additional single-cell assessment |
| GSE182227 | scRNA-seq | 4 patients | Oropharynx | p16 IHC-negative | Additional single-cell assessment |
| GSE208253 | Visium spatial transcriptomics | 10 patients; 12 sections | Oral cavity | Study-reported HPV-negative | Regional target–pEMT associations |
| GSE300414 | GeoMx spatial RNA profiling | 22 patients with paired tumor-bud and tumor-bulk regions | Oral cavity sites | p16 IHC-negative | Paired regional comparisons of cell states and target transcripts |
| CPTAC-HNSCC | Bulk RNA-seq and TMT proteomics | 108 patients; 84 with paired HER3 measurements | Oral cavity; larynx; oropharynx; lip; hypopharynx | Negative viral RNA inference<sup>c</sup> | RNA–protein concordance and cell-state associations |
| GSE290057 / GSE288406 | GeoMx RNA and antibody-signal profiling | 60 patients; one matched tumor region per patient | Pharynx; lip; tongue; oral cavity | Not reported | External RNA–antibody-signal agreement |
| S-BIAD3612 | CODEX multiplexed protein imaging | 28 cores; patient count unavailable | Larynx | Study-reported HPV-negative | Exploratory EGFR protein–state association |
| CCLE / DepMap | TMT proteomics and RNA-seq | 10 HNSCC cell lines; 8 complete for all three target proteins | Head and neck | Not reported | Exploratory protein–state associations in cell lines |
| TCGA-HNSC | Bulk RNA-seq and clinical data | 277 eligible patients; 273 with purity estimates | Oral cavity; larynx; pharynx | Specified HPV-assay eligibility rule | Bulk target-transcript balance and exploratory clinical associations |

**Abbreviations:** CCLE, Cancer Cell Line Encyclopedia; CPTAC, Clinical Proteomic Tumor Analysis Consortium; DepMap, Cancer Dependency Map; HNSCC, head and neck squamous cell carcinoma; HPV, human papillomavirus; IHC, immunohistochemistry; pEMT, partial epithelial–mesenchymal transition; scRNA-seq, single-cell RNA sequencing; TCGA, The Cancer Genome Atlas; TMT, tandem mass tag.

The discovery atlas included all sampled tissue contexts and HPV categories; within-patient malignant-cell analyses used eligible primary tumors. HPV ascertainment was retained as reported, distinguishing study-reported status, p16 immunohistochemistry and viral RNA inference. The primary analysis specification was timestamped locally before target–state testing, without public registration. The three pEMT between-target contrasts were primary in discovery and, following a documented amendment, in replication. Other analyses were exploratory. Cohorts were analyzed separately before exploratory synthesis. Patients, rather than cells, sections or regions, were the units of inference for patient cohorts^23^.

### Single-cell preprocessing, annotation and visualization

Analyses used deposited integer unique molecular identifier (UMI) counts, including counts extracted from author-filtered objects where applicable. Duplicate gene symbols were collapsed by summing counts. Quality control (QC) retained cells with more than 200 and fewer than 8,000 detected genes, at least 500 UMI and less than 10% mitochondrial UMI. Library totals comprised genes retained in each source matrix.

Discovery cell-type and malignant-cell annotations were retained from the original study. For GSE172577 and GSE188737, malignant-cell labels from the published non-immune HNSCC atlas were transferred by unique, exact within-patient matching of integer count vectors across shared genes^24^. GSE182227 used the original authors’ copy-number-alteration-based “Cancer” assignments; OP8 was excluded because of prior treatment, and HPV16 features were excluded from library totals^18^. GSE188737 patient HN279 was retained under the study-reported HPV-negative rule and excluded in a separate sensitivity analysis. Original counts, rather than integrated expression values, were used for target analyses.

Expression-inferred copy-number variation (CNV) and lineage-marker profiles provided annotation diagnostics. CNV analysis used infercnvpy with target-excluded autosomal genes and lineage-screened T-cell references. Discovery references were patient-matched when at least 20 qualified cells were available; otherwise, qualified cells from other patients were used. Cells whose mean squared CNV-window deviation exceeded the reference 95th percentile were designated CNV-supported. Original malignant labels defined the primary population; CNV-supported, lineage-consistent and joint restrictions were analyzed separately. External-cohort CNV diagnostics used the discovery parameters and lineage-screened T-cell references.

For display, counts were normalized to 10,000 UMI per cell and transformed as log(1 + CP10K). Embeddings used target-excluded highly variable genes, principal component analysis, Harmony, uniform manifold approximation and projection (UMAP), and Leiden clustering^25–27^. Nonmalignant cells in external-cohort atlases were labeled for display using discovery lineage-marker sets scored on Leiden clusters^27^. Embeddings were not used to estimate target effects, and expression maps were neither smoothed nor imputed. Transcript detection was defined as at least one observed UMI, with thresholds of two and three UMI examined as sensitivities. Detection fractions and mean normalized expression were calculated within patient-by-cell-type groups containing at least 10 cells and averaged with equal patient weight. Marker-expression colors were scaled within genes.

### Cell-state scores and within-patient target contrasts

The pEMT and two epithelial differentiation programs were obtained from Puram et al.. Gene symbols were reconciled using recorded mappings, and the three target genes were excluded from all programs^7^. The resulting sets contained 100, 99 and 99 genes, respectively; at least 70% of a program’s genes had to be available. A custom UCell-style score ranked positive expression values within each cell using descending average ranks^28^, assigning rank 1,501 to unexpressed genes and ranks beyond 1,500. For a program containing m available genes, the score was

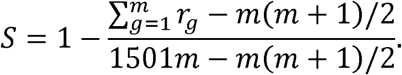

Patients required at least 50 quality-controlled malignant cells. Within each patient, the upper and lower thirds of the state-score distribution each contained floor(n/3) cells, with barcode order resolving ties deterministically. Comparisons required at least 10 cells per extreme, a mean score separation of at least 0.01, and at least 10 target UMI across the two extremes. For patient i and target g, the expression effect was

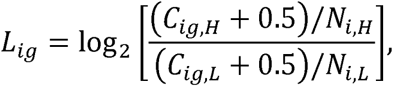

where C denotes summed target UMI and N denotes summed library UMI in the high (H) or low (L) group. The primary estimands were equal-patient means of the paired differences L(EGFR) − L(ERBB3), L(EGFR) − L(TACSTD2) and L(ERBB3) − L(TACSTD2). Each contrast included only patients with both target effects evaluable. These effects compare relative within-patient states rather than a common score increment across patients. The same procedure was applied to differentiation-1 and differentiation-2. Sensitivities varied the pseudocount to 0.1 or 1.0, the minimum malignant-cell count to 100 or 200, and the anatomical or cell-identity restrictions. For descriptive profiles, cells were divided into five approximately equal within-patient pEMT bins, with at least 10 cells per bin. Bin expression was log [(summed target UMI + 0.5)/summed library UMI × 10], centered across bins within each patient and target. Cohort profiles averaged these centered values with equal patient weight.

### Continuous associations and external-cohort synthesis

Within-patient partial Spearman correlations were calculated by regressing ranks of normalized target expression and pEMT score on ranked covariates and correlating the residuals. The technical-adjustment model included log library size and mitochondrial fraction; the differentiation-adjusted model additionally included differentiation-1 score. Between-target differences in partial correlations were calculated within patients before equal-patient aggregation. These analyses required estimable, nonconstant variables but did not impose the pseudobulk contrast’s minimum pooled target-UMI requirement.

Coupling between pEMT and differentiation-1 was summarized by within-patient Spearman correlations.

Continuous expression profiles were described using negative-binomial spline models of raw target counts in patients with at least 100 malignant cells. Models included a log library-size offset, patient intercepts and a cubic B-spline with four basis functions, with equal total weight assigned to each patient. Dispersion was estimated using an unweighted NB2 model before fitting the weighted negative-binomial generalized linear model. The descriptive pooled external model used within-cohort pEMT percentiles. Fitted log□-expression curves were centered over the prediction grid, and pointwise confidence bands were obtained by refitting models in patient-bootstrap samples. These analyses described cross-sectional variation along the fixed pEMT program, not temporal progression.

The exploratory external synthesis combined patient-level estimates from GSE172577, GSE188737 and GSE182227, excluding discovery. Patients received equal weight, and confidence intervals were obtained by resampling patients within cohorts while preserving cohort sample sizes. The same approach summarized the differentiation contrasts and pEMT–differentiation correlations. Replication in an individual additional cohort required both EGFR-relative pEMT contrasts to meet q < 0.05; pooled evidence was considered separately from this criterion.

### Visium spatial transcriptomics

GSE208253 comprised 12 sections from 10 patients with study-reported HPV-negative oral squamous cell carcinoma. Counts, spatial coordinates and registered hematoxylin and eosin images were linked to the authors’ pathologist annotations by barcode and coordinate agreement^19^. Spots required at least 500 UMI, at least 200 detected non-target genes and at most 20% mitochondrial UMI; spots labeled artifact, cautery, fold or edge effect were excluded. Primary analyses used pathologist-labeled SCC spots with composition proxy *f*_M_ ≥ 0.50.

The proxy *f*_M_ was the normalized malignant coefficient from nonnegative least-squares fitting to equal-donor discovery-atlas profiles, using CNV-supported malignant reference cells and reference-selected markers excluding targets, state-program genes and mitochondrial genes. Stromal and immune scores used DCN/LUM/FAP/PDGFRA and PTPRC/CD3D/CD79A/LST1, respectively. Rank-based scores used a cap of 819, selected without target-expression information from the 10th percentile of detected non-target genes across quality-controlled spots. Sections required at least 100 analyzed tumor spots and at least 20 nonzero spots per target.

Within each section, target expression was normalized to non-target UMI and transformed as log(1 + CP10K). Ranks of target expression and pEMT were residualized on ranks of *f*_M_, stromal score, immune score, log(1 + non-target UMI) and differentiation-1. Residual correlations defined partial Spearman estimates. The primary spatial contrasts were *Δ_H_ = p(EGFR) – p(ERBB3)* and *Δ_T_ = p(EGFR) – p(TACSTDD2*). Section estimates were averaged within patients before equal-patient aggregation. Section-level intervals used 2,000 spatial block-bootstrap resamples with 500-µm blocks; cohort-level intervals used 20,000 patient-bootstrap resamples.

Complementary analyses compared upper and lower within-section pEMT thirds using the pseudobulk estimand above. Sensitivities used *f*_M_ ≥ 0.70, all quality-controlled SCC spots, rank caps of 500 or 1,500, the five patients listed as receiving surgery alone, and leave-one-patient-out analyses. A post hoc adjustment ladder sequentially added *f*_M_ and depth, stromal and immune scores, and differentiation-1; the five-covariate model remained primary. Geometric interface spots were within 200 µm of pathologist-labeled stroma and more than 200 µm from tissue margins; core spots were at least 400 µm from stroma. Covariate-adjusted interface–core comparisons required at least 30 spots per region.

### GeoMx analysis of tumor buds and tumor-bulk regions

GSE300414 digital counts were reprocessed for paired PanCK-positive/CD45-negative tumor-bud and tumor-bulk areas of illumination (AOIs) from the same budding patients. Separate non-budding control patients were not used as paired comparators. Technical filters required at least 40,000 raw reads, stitched/raw and aligned/raw ratios of at least 0.80 and 0.70, respectively, and sequencing saturation of at least 0.50. The AOI-specific limit of quantification (LOQ) was the greater of 2 and the negative-probe geometric mean multiplied by the squared geometric standard deviation. Under the native convention, 0.5 was added to all negative counts in an AOI when any were below 0.5. AOIs required at least 5% of endogenous genes above LOQ, and genes required measurements above LOQ in at least 10% of retained AOIs.

The pipeline retained 215 AOIs and 6,844 genes overall, with paired tumor regions in 22 patients. Q3 factors were each AOI’s 75th-percentile count across retained genes divided by the geometric mean of these percentiles. Target signal was log [(raw count + 0.5)/Q3 factor]. AOI values were averaged within patient-region before calculating bud-minus-bulk differences and paired between-target contrasts. Observed values below LOQ were retained without imputation. Quantification was assessed separately: a target was above LOQ in a patient-region when at least one AOI exceeded its own threshold. Target-excluded pEMT and differentiation scores used the rank-sum formula with a rank cap of 1,500 within the retained gene universe and summarized using the same patient-region hierarchy. Sensitivities used zero-to-one negative-probe flooring with technical QC, or native negative probes with LOQ-only filtering^9,29^.

### CPTAC RNA–protein concordance and state associations

CPTAC-HNSCC RNA, protein and clinical data were matched by patient identifier. Primary analyses used the pan-cancer v1.2 data freeze: log□ upper-quartile-normalized RSEM RNA expression and reference-normalized log□ MS1 protein abundance from the tandem mass tag workflow^20,21^. Paired measurements were available for EGFR and TROP2 in 108 patients and HER3 in 84. Missing protein values were neither imputed nor classified as low expression.

RNA–protein concordance was quantified by Spearman correlation, with 95% percentile intervals from 5,000 patient-bootstrap samples and two-sided tests based on 19,999 patient-label permutations. Sensitivities adjusted for ranked whole-exome-sequencing-derived purity or used the original 2021 processing of the same cohort. Between-target differences in concordance used the 84 patients complete for all three targets and 5,000 shared patient-bootstrap samples, preserving pairing across targets.

Bulk pEMT and differentiation scores used the target-excluded rank-sum approach with a rank cap of 1,500 within the downloaded 60,669-gene RNA-expression universe. RNA–state and protein–state correlations were compared in the same complete patient set for each target, with identical patient sets used for pEMT and differentiation-1: 108 patients for EGFR and TROP2 and 84 for HER3. RNA-minus-protein differences in Spearman correlations were evaluated by shared patient bootstrap, resampling RNA, protein and state scores jointly. Separate available-case analyses of both differentiation programs retained all available measurements within each modality.

Exploratory high-expression groups were defined separately for RNA and protein using the linearly interpolated 75th percentile, retaining boundary ties. Agreement was the proportion of RNA-upper-quartile cases also in the protein upper quartile; non-overlap was its complement. The principal comparison used the 84 complete cases. Intervals used 20,000 shared patient-bootstrap samples, recalculating thresholds and group sizes in each sample. Upper-third and upper-half definitions, target-specific available cases and original processing were sensitivities. These groups represented cohort-relative rankings rather than clinical cutoffs.

### Complementary protein analyses

External assay agreement used 60 patients with one tumor region per patient matched between GeoMx RNA and antibody-signal assays on serial sections (GSE290057/GSE288406). Deposited relative-log-expression-normalized log□ counts per million were analyzed using the same rank-group definitions. Target antibody signals were compared with the maximum of five IgG controls in each region as a background diagnostic, not a validated LOQ^30^.

CODEX analyses used deposited per-cell whole-cell intensities and Nimbus scores from 28 laryngeal SCC cores in S-BIAD3612. Patient identifiers were unavailable, so each core was assumed to represent one patient. Tumor-cell selection used PanCK or TP63 positivity, exclusion of CD8- or CD68-positive cells, and available tumor boundaries; target and state-proxy markers were not used for gating. Each core required at least 100 eligible cells. The state proxy was the mean within-core percentile rank of LAMC2 and podoplanin. Partial Spearman correlations with EGFR and control markers, including ATP1A1 and cytochrome c, were adjusted for log cell area and DAPI intensity. Intensity- and Nimbus-based analyses were performed separately. Core-level means used 20,000 bootstrap resamples and 99,999 Monte Carlo sign flips^31^.

CCLE tandem mass tag proteomics was paired with DepMap 24Q4 log (TPM + 1) RNA data. RNA state scores were means of within-population gene-wise z scores for available target-excluded signature genes. Protein associations used individual targets and the balance index z(EGFR) − ½[z(HER3) + z(TROP2)], without imputing missing protein. Analyses included 10 HNSCC lines, eight protein-complete, and a lineage-adjusted sensitivity across 34 squamous lines, 28 protein-complete^32–34^. Inference treated cell lines as units, using permutation tests and 20,000 line-bootstrap samples; the broader analysis preserved lineage strata during resampling.

### TCGA molecular and clinical analyses

TCGA-HNSC HiSeqV2 log (RSEM + 1) expression, clinical annotations, GISTIC2 gene-level copy number and ABSOLUTE purity were obtained from UCSC Xena and PanCanAtlas resources. Overall survival (OS) and disease-specific survival (DSS) were obtained from the TCGA Clinical Data Resource^22,35–38^. One primary tumor per patient was retained. Primary eligibility required at least one negative result and no positive result among p16 immunohistochemistry, HPV in situ hybridization and the TCGA RNA-seq HPV call. MassArray HPV-DNA status was not an eligibility assay. A broader sensitivity population included known non-oropharyngeal tumors without a positive result on the specified assays, including cases with unknown HPV status.

Expression was standardized within each eligibility population. Bulk program scores were means of gene-wise z scores for available target-excluded signature genes. Target-transcript balance was defined as

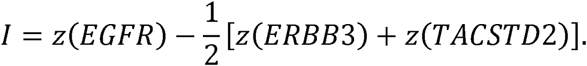

Partial Spearman correlations of individual targets and I with pEMT were adjusted for ABSOLUTE purity. Additional models included a mean-z-score proxy of ESTIMATE stromal genes or target-specific copy-number values; the index model included copy number for all three targets^39^. Post hoc eligibility sensitivities excluded MassArray-positive cases or restricted analyses to RNA-seq HPV-negative cases.

Pathological nodal involvement (N1–N3 versus N0) was modeled by logistic regression adjusted for age, sex, anatomical subsite, smoking, pathological T category and purity. Cox models for OS and DSS adjusted for age, pathological stage, subsite, sex, smoking and purity. Complete-case analysis was used, and odds ratios and hazard ratios were expressed per one-standard-deviation increase in I within the eligibility population. Proportional hazards were assessed using Schoenfeld-residual-based tests with rank-transformed time; subsite-stratified Cox models were post hoc sensitivities. Kaplan–Meier displays used a median index split and were descriptive, separate from the continuous-index regression models^40^.

### Synthetic pseudobulk analysis

Synthetic expected-count pseudobulks were constructed from seven discovery patients with at least 50 malignant cells, 20 fibroblasts and 20 immune cells. Patient-specific mean raw-count profiles were calculated for pEMT-high and pEMT-low malignant thirds, fibroblasts and immune cells. Mixing weights represented cell-number fractions, retaining differences in mean UMI content between compartments.

Four designs were evaluated. C1 varied malignant-cell fraction from 0.3 to 0.9 while holding the malignant high:low ratio at 1:1 and dividing the remainder equally between fibroblasts and immune cells. C2 fixed malignant fraction at 0.5 and varied fibroblast fraction from 0.05 to 0.45. S fixed malignant, fibroblast and immune fractions at 0.70, 0.15 and 0.15, respectively, while varying the pEMT-high share of malignant cells from 0.1 to 0.9. M varied the same state share in malignant-only mixtures.

Mixtures were normalized as log (CPM + 1). Synthetic pEMT was the mean standardized expression of available program genes, using a common scale derived from the deterministic mixtures. Within-patient slopes related the EGFR−ERBB3 and EGFR−TACSTD2 expression balances to synthetic pEMT. A further 1,000 composition-only cohorts contained one random mixture per patient, with the malignant high:low ratio fixed at 1:1. Between-patient Spearman correlations were summarized by their median and 2.5th–97.5th percentiles. These intervals describe simulation variability conditional on the source patients, not confidence intervals for an observed tumor population.

### Statistical analysis

Mean patient-level effects were tested using two-sided sign-flip tests, enumerating all sign patterns for up to 23 independent units and using 99,999 Monte Carlo flips with a plus-one correction above that number. Tests assumed independent units and symmetry of effects under the null^41,42^. Comparisons with fewer than two evaluable units were descriptive and were not tested. Single-cell and spatial cohort-level mean effects used 20,000 percentile-bootstrap samples; pooled external estimates resampled patients within cohorts. Spatial section-level, spline, concordance and assay-group analyses used their analysis-specific resampling procedures. TCGA partial-correlation intervals used Fisher-z approximations, and logistic and Cox estimates used Wald intervals^43^. Confidence intervals were pointwise, not simultaneous or multiplicity-adjusted.

Benjamini–Hochberg correction was applied within defined hypothesis families. The single-cell pEMT family comprised three between-target contrasts per cohort; the six contrasts across differentiation-1 and differentiation-2 formed a separate family. The corresponding three- and six-contrast families were used for the pooled external synthesis. Differentiation-adjusted individual-target correlations and EGFR-minus-partner differences formed separate three- and two-test families. Visium between-target contrasts, individual-target correlations, pseudobulk contrasts and interface–core comparisons were adjusted separately. GeoMx target effects, between-target contrasts and state-program differences each formed separate three-test families. CPTAC concordance used a three-target family; TCGA molecular families comprised the three targets and I, and its clinical family comprised nodal involvement, OS and DSS. Adjustment used finite, testable p values, with q < 0.05 defining significance within each family^44^. Exploratory paired correlation differences and rank-group agreement were summarized by pointwise intervals; simulations were descriptive. Post hoc TCGA sensitivity models were reported with nominal p values.

### Software, reproducibility and AI assistance

Analyses were implemented in Python using NumPy, SciPy, pandas, Scanpy, AnnData, infercnvpy, harmonypy, scikit-learn, statsmodels, patsy and lifelines^45^. Main figures were prepared in R with ggplot2 and associated layout packages from exported analysis tables, without recomputing estimates during plotting; most supplementary figures were prepared with Matplotlib. Pinned software versions, analysis-specific random seeds, patient-level inclusion records are retained in the reproducible code archive. AI agents (OpenAI Codex and Anthropic Claude) assisted with code development, manuscript language polishing. Scientific plots were generated programmatically from analysis outputs; no generative image synthesis was used.

## Results

### A single-cell atlas maps the cellular distribution of EGFR, ERBB3 and TACSTD2 transcripts

The discovery atlas comprised 54,239 quality-controlled cells from 37 samples collected from 23 patients in GSE181919 (Fig. 1A; Supplementary Fig. S1A–G). Primary tumors, adjacent mucosa, leukoplakia and lymph-node metastases represented the sampled tissue contexts, with reported HPV-positive and HPV-negative cases included in the atlas (Supplementary Fig. S1B–D). Lineage-marker expression supported the major epithelial, stromal and immune populations, providing a cellular framework for examining the three target transcripts (Fig. 1B; Supplementary Fig. S2A).

**Figure 1.**
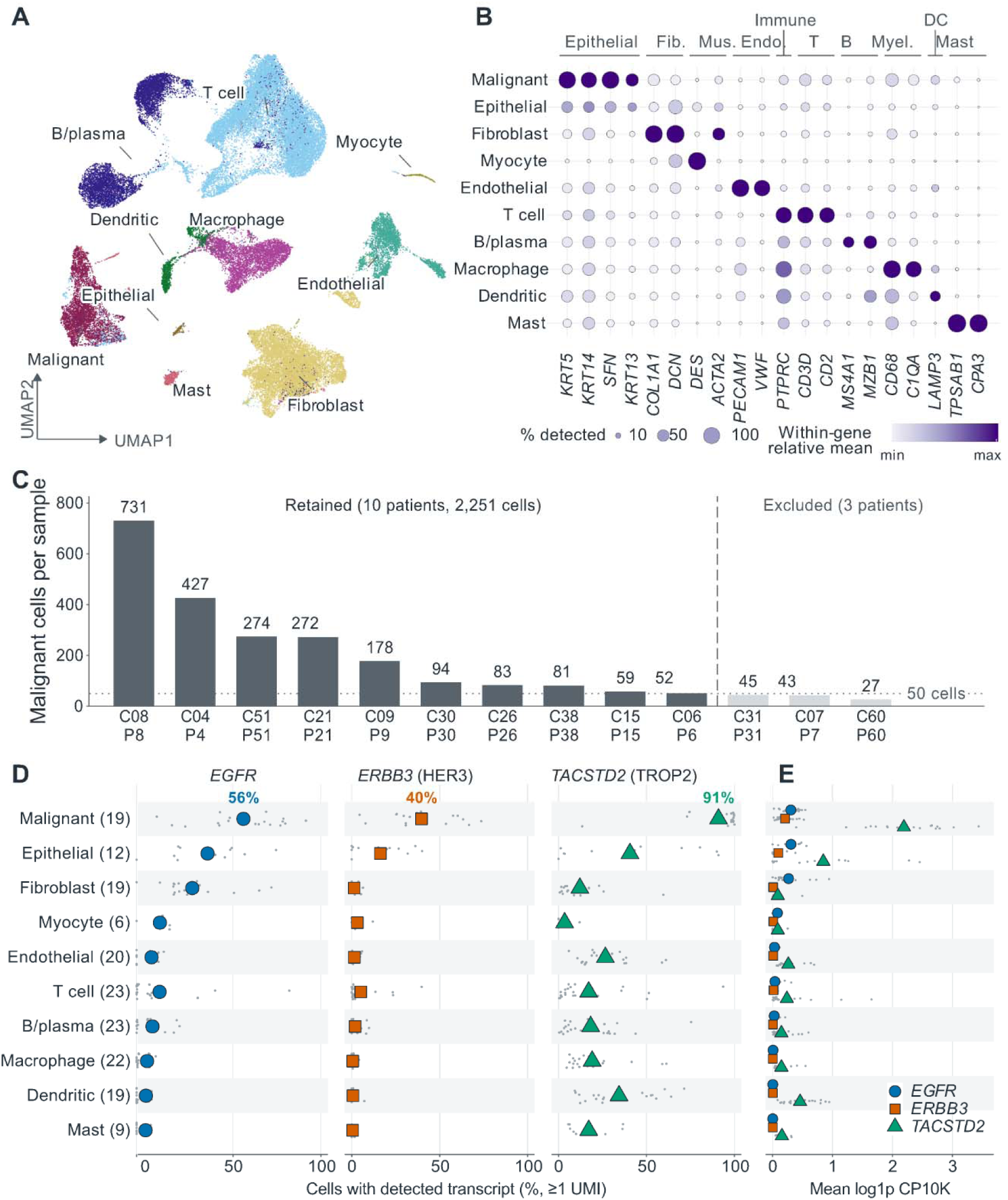
Cellular distribution of EGFR, ERBB3 and TACSTD2 transcripts in the discovery single-cell atlas. (A) UMAP of 54,239 cells from 37 samples and 23 patients (GSE181919), colored by source cell-type annotations. (B) Lineage-marker expression across cell types. Dot size indicates equal-patient mean detection; color indicates within-gene scaled mean normalized expression. (C) Malignant-cell counts in reported HPV-negative primary tumors (n = 13), showing the 10 patients retained (> 50 cells; 2,251 cells, dark bars) and three excluded (< 50 cells, light bars) for within-patient analyses. (D,E) Cellular detection fractions and mean normalized expression for EGFR (blue *circles), ERBB3 (HER3, orange squares) and TACSTD2 (TROP2, green triangles). Small gray points show individual patients and large symbols show equal-patient means; percentages in D mark malignant-cell means*.

Primary tumors reported as HPV-negative contributed 2,366 cells annotated as malignant from 13 patients. Expression-inferred copy-number profiles supported these annotations in at least 89% of cells in 11 of the 13 patients, with patient-level fractions ranging from 59% to 100% (Supplementary Fig. S2B–D). Ten patients met the requirement of at least 50 malignant cells, yielding 2,251 cells for within-patient analyses (Fig. 1C).

The three targets differed in their cellular distribution. Across the full atlas, TACSTD2 was the most broadly detected transcript in malignant cells: among the 19 contributing patients, equal-patient mean detection fractions were 91% for TACSTD2, 56% for EGFR and 40% for ERBB3 (≥1 UMI; Fig. 1D). ERBB3 detection was concentrated in epithelial compartments, whereas EGFR was also detected in fibroblasts and TACSTD2 in endothelial cells and several immune populations. TACSTD2 showed its highest patient-averaged normalized expression in malignant cells (Fig. 1E).

Broad malignant-cell detection of TACSTD2 persisted at thresholds of at least two or three UMIs, while the lower-count signals in non-epithelial populations were more threshold-sensitive (Supplementary Fig. S3A). The ordering of malignant-cell detection fractions was retained in the primary-tumor subset used for within-patient analyses (Supplementary Fig. S3B).

### Target-transcript profiles shift toward EGFR across intratumoral pEMT states

We next examined how target expression varied across partial epithelial–mesenchymal transition (pEMT) states within individual tumors. Target-excluded pEMT and epithelial differentiation-1 scores varied across discovery-cohort malignant cells, alongside heterogeneous EGFR, ERBB3 and TACSTD2 expression (Fig. 2A,B). Patient-level analyses included 10 discovery patients, six replication patients (GSE172577), six patients from GSE188737 and four from GSE182227, each with at least 50 malignant cells (Fig. 2C). Cohort-specific atlases, marker profiles and copy-number analyses provided the corresponding cellular and annotation context (Supplementary Figs. S4A–F and S5A,B).

**Figure 2.**
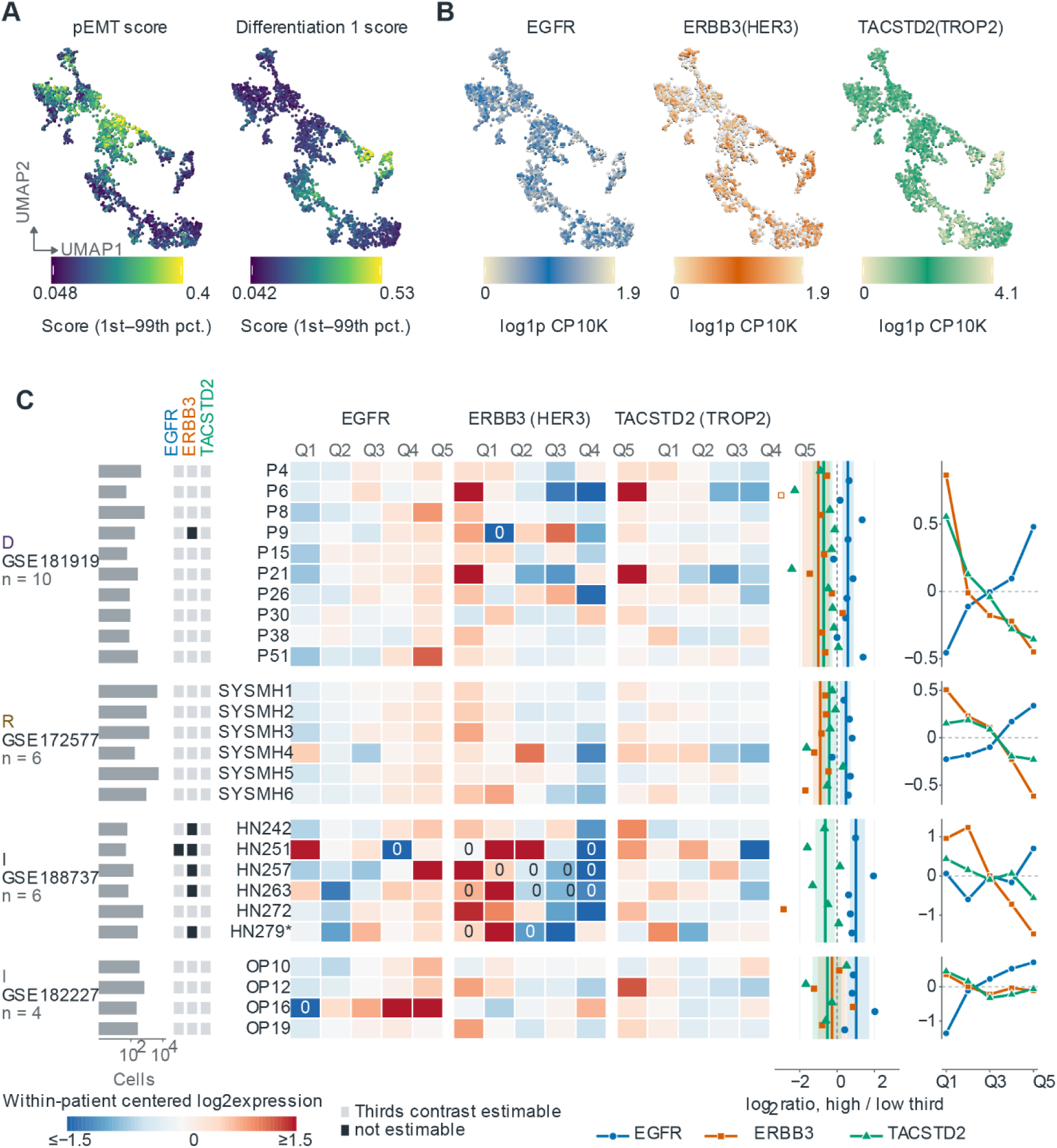
Malignant-cell states and target-transcript expression across within-patient pEMT strata. (A) UMAP of discovery-cohort malignant cells colored by pEMT and epithelial differentiation scores (calculated with target genes excluded). (B) The same UMAP colored by target-transcript expression (log p CP10K, unsmoothed). (C) Target expression across within-patient pEMT quintiles (Q1–Q5). The leftmost bars indicate patient-specific malignant-cell counts across four cohorts (discovery, replication, and *independent cohorts). Adjacent gray columns indicate estimable (light) or non-estimable (dark) effects for each target. Heatmaps show patient-centered target expression across quintiles; a printed “0” indicates no observed target UMI. Right panels show patient-level log ratios of expression in the upper versus lower pEMT thirds (dots) and cohort-level mean profiles (lines). Blue circles, orange squares, and green triangles denote EGFR, ERBB3, and TACSTD2, respectively. The asterisk marks patient HN279 (p16-positive)*.

Within-patient pEMT quintiles (Q1, lowest; Q5, highest) revealed contrasting target-expression profiles. EGFR expression was higher in Q5 than in Q1 in all 10 discovery patients and five of six replication patients. The corresponding counts were one of 10 and none of six for ERBB3, and one of 10 and two of six for TACSTD2 (Fig. 2C). Equal-patient means of the within-patient-centered log-expression profiles showed higher EGFR and lower ERBB3 and TACSTD2 expression at the upper end of the pEMT distribution in both cohorts (Fig. 2C). Patient-specific quintile positions and raw target-count support are shown in Supplementary Fig. S6A,B.

Comparisons of the upper and lower pEMT thirds quantified these patterns. In discovery tumors, the mean log high-to-low expression ratio was 0.58 for EGFR (95% CI, 0.28–0.89; n = 10), −1.00 for ERBB3 (95% CI, −1.87 to −0.40; n = 9) and −0.72 for TACSTD2 (95% CI, −1.30 to −0.25; n = 10). Replication-cohort means were 0.48, −0.90 and −0.43, respectively, with all six patients evaluable for each target (Fig. 2C).

The two additional cohorts extended the EGFR–TACSTD2 pattern. EGFR expression was higher in the pEMT-high third in all five evaluable GSE188737 patients and all four GSE182227 patients, with mean log ratios of 1.01 and 1.03. Mean TACSTD2 effects were −0.64 and −0.51, respectively. ERBB3 effects were evaluable in one of six GSE188737 patients and all four GSE182227 patients, with variable directions in the latter cohort (Fig. 2C;). Thus, the positive mean EGFR and negative mean TACSTD2 effects recurred across all four cohorts.

The same directions were evident in descriptive models along the continuous pEMT score. In patients with at least 100 malignant cells, fitted expression was higher for EGFR and lower for ERBB3 and TACSTD2 at the upper than at the lower end of the score range in discovery tumors and the pooled external cohorts (Supplementary Fig. S7A,B). Cohort-specific profiles show the variation around these aggregate patterns (Supplementary Fig. S7C–E).

### Direct within-patient comparisons distinguish EGFR from ERBB3 and TACSTD2

Paired comparisons showed a more positive within-patient pEMT association for EGFR than for ERBB3 or TACSTD2. In discovery tumors, the mean difference between high-to-low log□ expression ratios was 1.58 for EGFR−ERBB3 (95% CI, 0.87–2.42; n = 9) and 1.30 for EGFR−TACSTD2 (95% CI, 0.75–1.91; n = 10), with both contrasts positive in every evaluable patient (both q = 0.006). The corresponding replication estimates were 1.39 (95% CI, 1.06–1.80) and 0.91 (95% CI, 0.63–1.17), again positive in all six patients (both q = 0.047; Fig. 3A).

**Figure 3.**
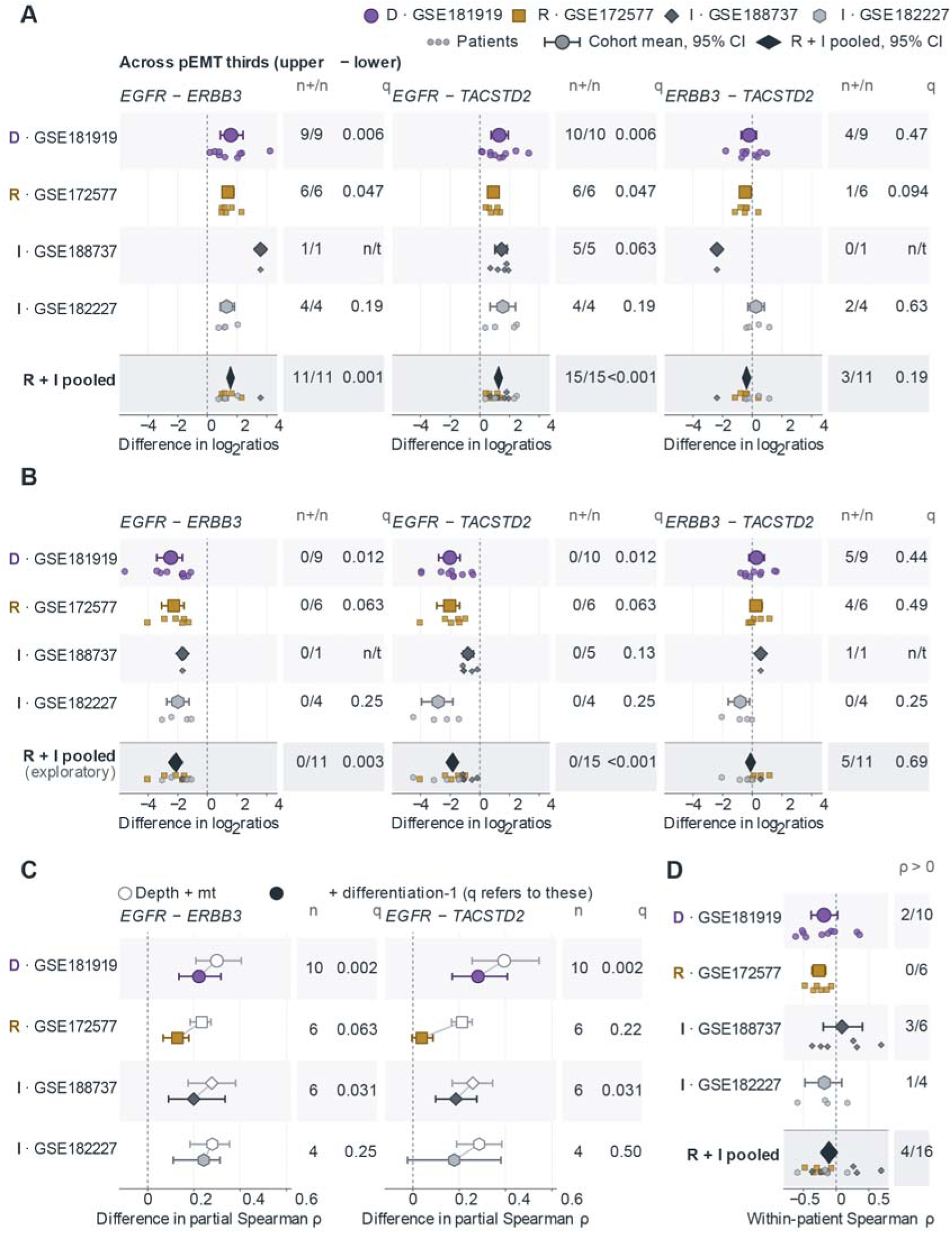
Relative target-transcript associations with pEMT and epithelial differentiation across single-cell cohorts. (A) Paired differences in target-specific log expression ratios between upper and lower within-patient pEMT thirds. Positive values indicate a stronger pEMT-associated difference for the first-named target. (B) Paired between-target differences comparing upper versus lower differentiation-1 thirds. Negative values indicate a weaker differentiation-associated difference for the first-named target. (C) Paired differences in partial Spearman correlations of target expression with pEMT. *Open symbols are adjusted for library size and mitochondrial fraction; filled symbols are further adjusted for differentiation-1. (D) Within-patient Spearman correlations between pEMT and differentiation-1 scores*.

The evaluable EGFR contrasts in both additional cohorts were directionally concordant. In an exploratory synthesis of the replication and two additional cohorts, excluding discovery, the EGFR−ERBB3 and EGFR−TACSTD2 contrasts were positive in all 11 and all 15 evaluable patients, respectively (q = 0.001 and q < 0.001). The ERBB3−TACSTD2 comparison did not support a consistent ordering of these two targets after multiplicity adjustment (Fig. 3A). Excluding the p16-positive patient HN279, retained under the study-reported HPV-negative eligibility rule, left positive EGFR−TACSTD2 contrasts in all four remaining evaluable GSE188737 patients (mean, 1.65 log units; 95% CI, 1.35–1.87; q = 0.125).

Epithelial differentiation-1 showed the opposite relative target pattern. Across differentiation-1 thirds, both EGFR-relative contrasts were negative in every evaluable patient in all four cohorts, indicating more positive differentiation-associated expression differences for ERBB3 and TACSTD2 than for EGFR. Discovery means were −2.47 and −2.03 log units (both q = 0.012); both contrasts were also negative in all six replication patients (both q = 0.063; Fig. 3B).

In the exploratory pooled external differentiation analysis, the EGFR−ERBB3 contrast was negative in all 11 evaluable patients (mean, −2.12 log units; 95% CI, −2.67 to −1.64; q = 0.003), and the EGFR−TACSTD2 contrast was negative in all 15 (mean, −1.83; 95% CI, −2.46 to −1.26; q < 0.001). These q values account for all six between-target contrasts across differentiation programs 1 and 2 (Fig. 3B).

Within-patient pEMT and differentiation-1 scores were negatively correlated in eight of 10 discovery tumors and all six replication tumors, with mean Spearman correlations of −0.19 and −0.26. Relationships were more variable in the additional cohorts (Fig. 3D). We next assessed how differentiation adjustment affected continuous target–pEMT associations. After adjustment for library depth, mitochondrial fraction and differentiation-1 score, the EGFR−ERBB3 difference in partial Spearman ρ remained positive in every discovery and replication patient, with means of 0.222 and 0.129 (q = 0.002 and q = 0.063, respectively). Both adjusted target contrasts were also positive in all six GSE188737 patients (both q = 0.031; Fig. 3C).

The EGFR−TACSTD2 contrast showed greater overlap with differentiation, particularly in replication tumors. Adding differentiation-1 score reduced its mean partial-correlation difference from 0.394 to 0.281 in discovery (adjusted 95% CI, 0.170–0.408; q = 0.002) and from 0.213 to 0.038 in replication (95% CI, −0.004 to 0.086; q = 0.219). The adjusted GSE182227 estimate was also imprecise (q = 0.50; Fig. 3C). Together, the unadjusted contrasts consistently separated EGFR from both partner targets, whereas differentiation adjustment attenuated the EGFR−TACSTD2 separation, particularly in replication tumors.

### Spatial profiling supports the relative EGFR–pEMT association within tumor regions

We examined the regional counterpart of the single-cell findings in 12 Visium sections from 10 patients with study-reported HPV-negative oral SCC (GSE208253). The primary analysis comprised 14,833 quality-controlled spots within pathologist-annotated SCC regions meeting the composition-proxy threshold fM ≥ 0.50 (Fig. 4A,B; Supplementary Fig. S8A,B). Associations were estimated within sections after adjustment for composition proxies, library depth and differentiation-1 score, then averaged within patient.

**Figure 4.**
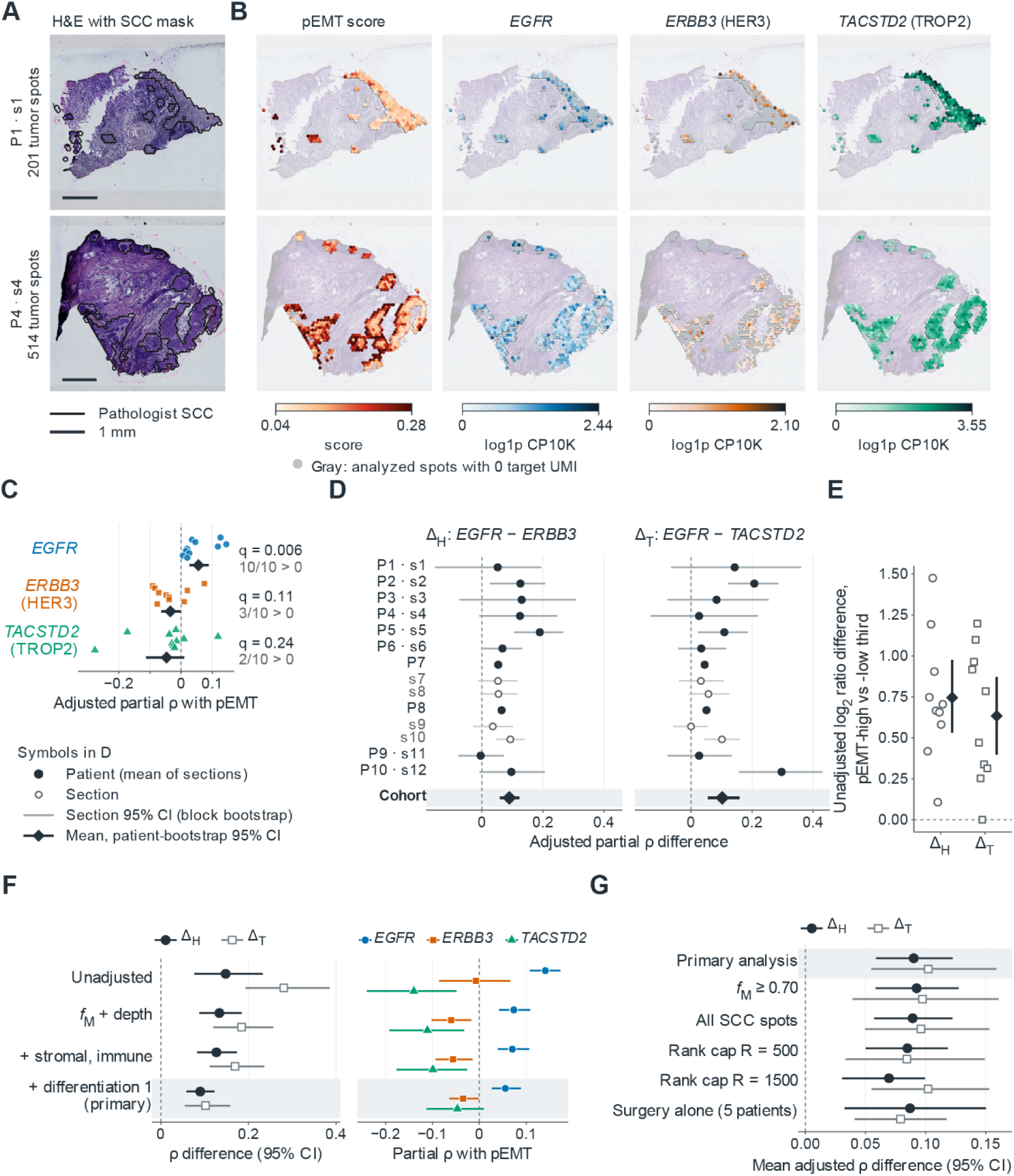
Spatial associations of EGFR, ERBB3, and TACSTD2 transcripts with pEMT in oral squamous cell carcinoma. Spatial transcriptomics data from 12 Visium sections across 10 patients (GSE208253). (A) H&E images with the pathologist-annotated SCC mask for two representative patients. (B) Unsmoothed spatial maps of pEMT score and *target expression. Gray indicates spots with zero target UMI. (C) Within-section adjusted partial correlations of each target with pEMT. (D) Primary differences in correlations:* Δ*H (EGFR – ERBB3) and* Δ*T(EGFR – TACSTD2). (E) Unadjusted pseudobulk log expression ratio differences (pEMT-high vs. pEMT-low thirds). (F) Post hoc adjustment ladder for correlation differences (left) and individual partial correlations (right). The locked primary model is shaded. (G) Sensitivity analysis showing* Δ*H and* Δ*T across different composition and scoring settings. In C–G, small symbols represent individual sections/patients, and large diamonds with bars indicate equal-patient means with 95% confidence intervals*.

EGFR showed a more positive regional association with pEMT than either partner target. The mean paired difference in partial Spearman ρ was 0.090 for EGFR−ERBB3 (95% CI, 0.059–0.123; nine of 10 patients positive) and 0.102 for EGFR−TACSTD2 (95% CI, 0.055–0.159; all 10 positive; both q = 0.004; Fig. 4D). EGFR itself had a modest positive correlation with pEMT (mean partial ρ, 0.056; 95% CI, 0.027–0.089; q = 0.006), whereas the ERBB3 and TACSTD2 mean correlations were negative but did not individually meet the multiplicity-adjusted significance threshold (q = 0.111 and q = 0.240; Fig. 4C).

The relative ordering was consistent across complementary analyses. Unadjusted pseudobulk comparisons of the upper and lower within-section pEMT thirds gave positive EGFR−ERBB3 and EGFR−TACSTD2 contrasts in all 10 patients, with mean differences of 0.75 and 0.63 log units (both q = 0.002; Fig. 4E). In a post hoc adjustment ladder, the mean between-target differences decreased as covariates were added but remained positive after adjustment for composition proxies, depth and differentiation-1 (Fig. 4F). Mean adjusted differences also remained positive under alternative composition thresholds, inclusion of all quality-controlled SCC spots, alternative scoring rank caps and restriction to the five patients listed as receiving surgery alone (Fig. 4G). Patient-level and leave-one-patient-out summaries supported the same direction (Supplementary Fig. S9D,E).

The separate geometric interface–core comparison showed no clear pEMT or target-expression separation in the eight eligible patients. The composition proxy also provided limited separation of SCC and non-SCC regions (Supplementary Fig. S9A–C). These estimates describe associations among multicellular spots. Within that scope, the Visium analyses support the relative EGFR–pEMT pattern observed in the single-cell cohorts.

### Tumor buds combine higher pEMT with lower epithelial differentiation

To relate the pEMT and differentiation patterns to histologically defined compartments, we compared tumor buds with matched tumor-bulk regions. In GSE300414, 22 patients with negative p16 immunohistochemistry contributed paired, quality-controlled PanCK-positive/CD45-negative tumor regions, comprising 38 bud and 43 tumor-bulk areas of illumination (AOIs; Supplementary Fig. S10A–C). Here, tumor bulk denotes a sampled region within the same budding case.

Tumor buds had a pEMT-high, less differentiated transcriptional profile. pEMT scores were higher in buds in 20 of 22 patients, with a mean bud-minus-bulk difference of 0.110 (95% CI, 0.076–0.146). Differentiation-1 scores were lower in buds in 20 of 22 patients (mean difference, −0.162; 95% CI, −0.220 to −0.110), and differentiation-2 scores were lower in 21 of 22 (mean difference, −0.059; 95% CI, −0.076 to −0.041). All three exploratory state comparisons had q < 0.0001 (Fig. 5A–C).

**Figure 5.**
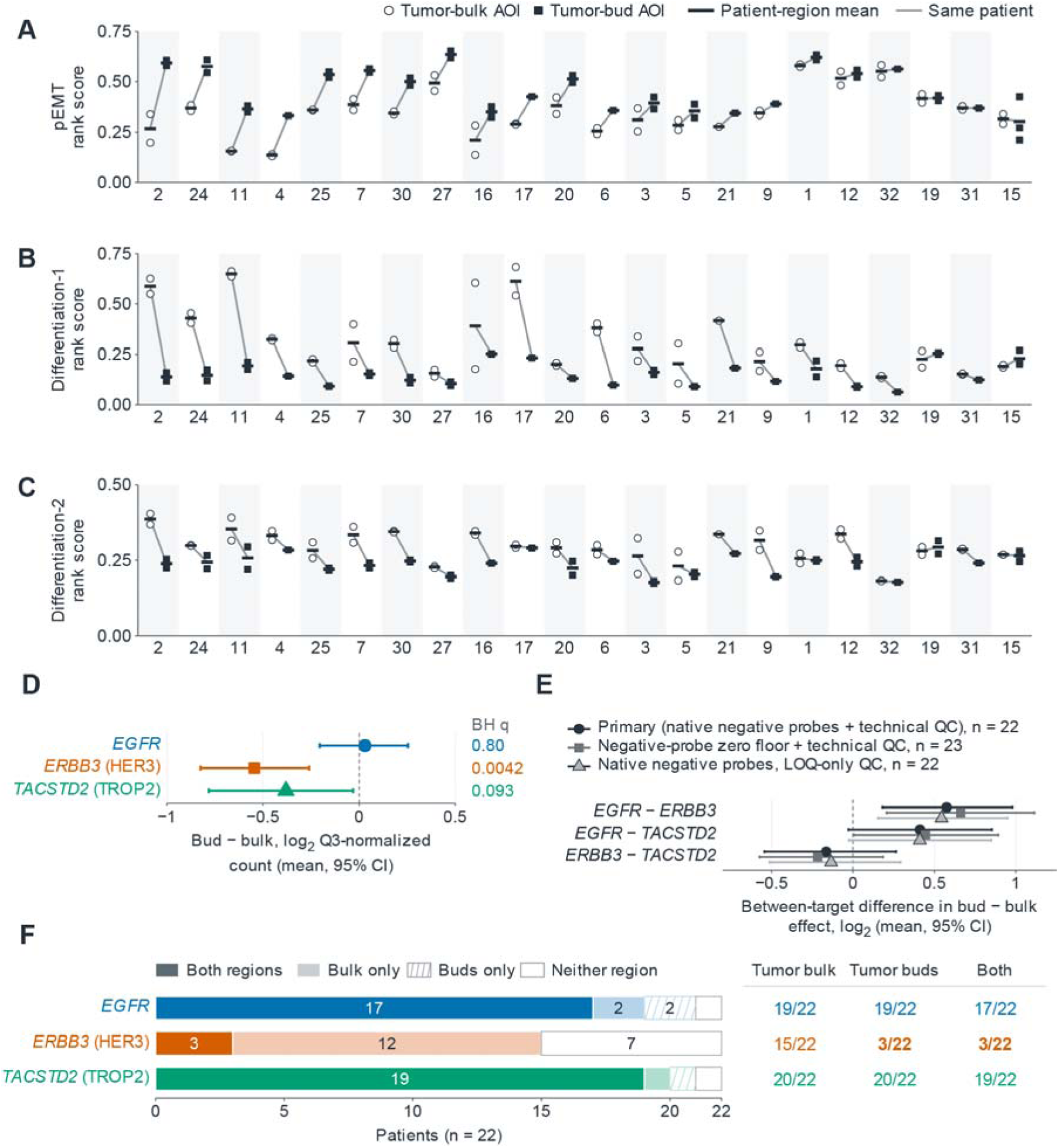
Tumor buds are high-pEMT, low-differentiation regions. Spatial transcriptomics (GeoMx) profiling of tumor bulk and tumor bud regions from 22 HNSCC patients (GSE300414). (A–C) Rank scores for pEMT (A), differentiation-1 (B), and differentiation-2 (C) per region of interest (AOI). Circles indicate tumor bulk; squares indicate tumor buds; lines connect AOIs from the same patient. (D) Differences in normalized target expression between tumor buds and bulk for EGFR, ERBB3, and TACSTD2. (E) Between-target expression contrasts across different quality-control (QC) conventions. (F) Summary of patients with target expression above the limit of quantification (LOQ) across bulk, bud, or both regions. In A–E, points/bars represent means with 95% confidence intervals.

Analyses retaining all observed counts, including measurements below the limit of quantification (LOQ), showed target-specific regional signal differences. The mean bud-minus-bulk difference in log□ Q3-normalized signal was close to zero for EGFR (0.03; 95% CI, −0.21 to 0.25; q = 0.80), negative for ERBB3 (−0.54; 95% CI, −0.83 to −0.26; q = 0.0042), and negative for TACSTD2 without meeting the multiplicity-adjusted threshold (−0.38; 95% CI, −0.78 to −0.03; q = 0.093; Fig. 5D). The EGFR−ERBB3 contrast was 0.58 log units (95% CI, 0.18–0.98; q = 0.035), and the EGFR−TACSTD2 contrast was 0.41 (95% CI, −0.03 to 0.85; q = 0.135; Fig. 5E). Both mean contrasts remained positive under alternative QC conventions (Fig. 5E).

EGFR and TACSTD2 measurements exceeded the AOI-specific LOQ in at least one AOI from each region in 17 and 19 of 22 patients, respectively. ERBB3 had an above-LOQ measurement in at least one tumor-bulk AOI in 15 patients and in at least one bud AOI in three patients (Fig. 5F). At the AOI level, four of 38 bud measurements and 23 of 43 bulk measurements exceeded the ERBB3 LOQ (Supplementary Fig. S10D); regional mean ERBB3 count/LOQ ratios were below 1 in 20 of 22 bud regions.

Because most bud ERBB3 measurements were below LOQ, its contrast describes an all-count signal difference rather than a precisely quantified regional decrease. These paired regional data characterize tumor buds as a pEMT-high, less differentiated compartment with target-specific transcript-signal profiles.

### Paired RNA and protein measurements reveal target-specific concordance and state associations

Paired measurements in CPTAC-HNSCC showed positive RNA–protein correlations for all three targets, with particularly close correspondence for EGFR and TROP2. Spearman correlations were 0.862 for EGFR RNA and EGFR protein (95% CI, 0.795–0.906; n = 108), 0.764 for TACSTD2 RNA and TROP2 protein (95% CI, 0.678–0.824; n = 108), and 0.404 for ERBB3 RNA and HER3 protein (95% CI, 0.198–0.578; n = 84; Fig. 6A–C). The ordering was retained after adjustment for whole-exome sequencing-derived tumor purity and in the original 2021 processing of the same cohort (Fig. 6D). In exploratory paired comparisons of the 84 complete cases, the RNA–protein correlation was higher for EGFR than for HER3 by 0.460 (95% CI, 0.261–0.663) and higher for TROP2 than for HER3 by 0.335 (95% CI, 0.125–0.537; Fig. 6E).

**Figure 6.**
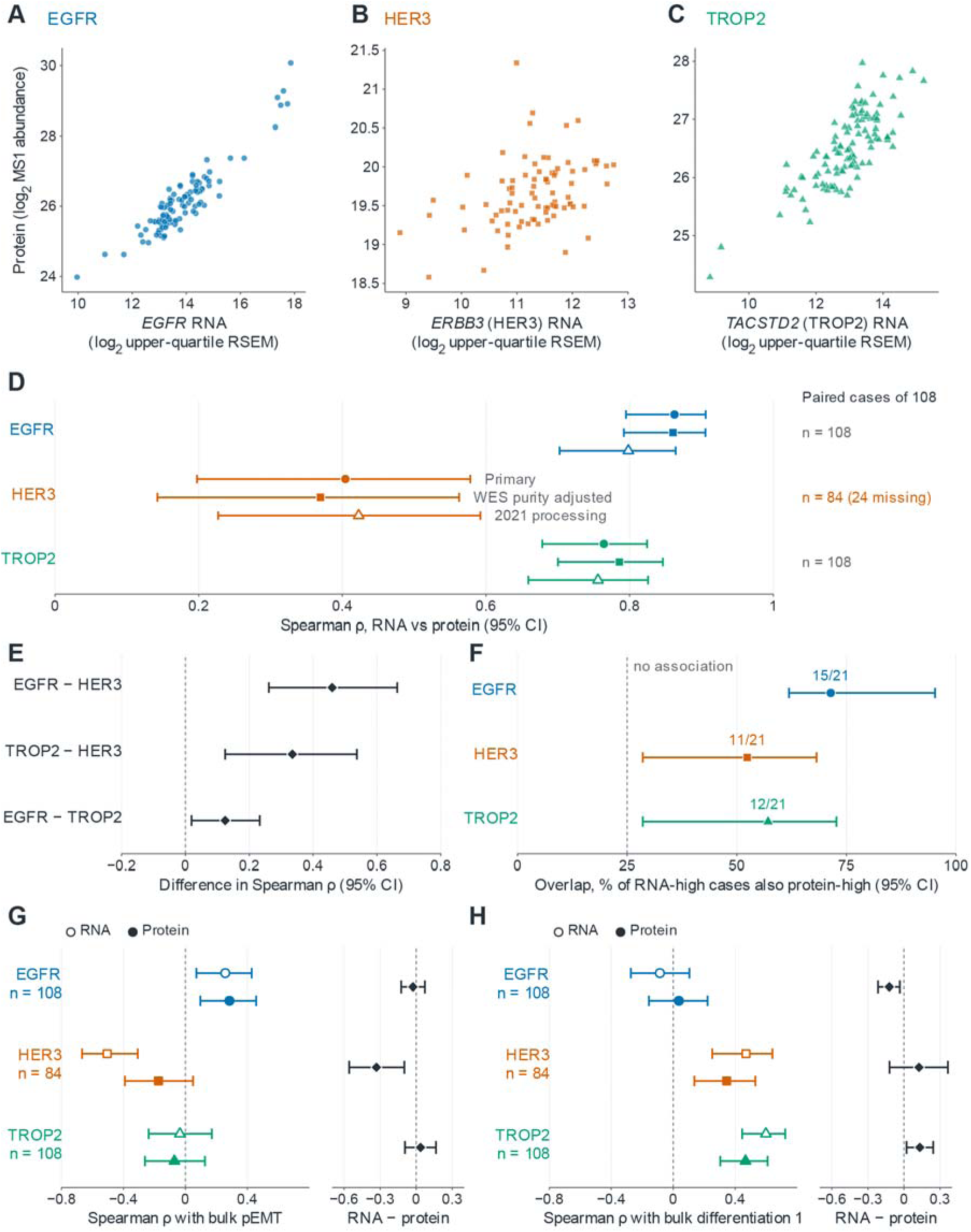
Target-specific RNA–protein concordance and their associations with pEMT and epithelial differentiation in CPTAC-HNSCC. Analyses of primary tumors from CPTAC-HNSCC. (A–C) Correlation between paired RNA and protein abundance for EGFR, HER3, and TROP2. (D) Spearman correlations, showing primary estimates alongside those adjusted for tumor purity and processing. (E) Paired differences in RNA–protein correlations between targets. (F) Overlap of RNA and protein high-expression *cases. (G, H) Correlations of target RNA (open symbols) and protein (filled symbols) with bulk pEMT (G) and differentiation-1 (H) scores. Right panels show paired RNA minus protein correlation differences. Bars indicate 95% confidence intervals*.

Exploratory upper-quartile comparisons used the 84 cases complete for all three targets. Of 21 RNA-upper-quartile patients per target, 15 were also in the EGFR protein upper quartile (71.4%), 11 in the HER3 upper quartile (52.4%) and 12 in the TROP2 upper quartile (57.1%; Fig. 6F; Supplementary Fig. S11A–C). Complementary non-overlap estimates and upper-third and upper-half comparisons are provided in Supplementary Fig. S11D–F. These groups describe within-cohort rankings rather than clinical positivity thresholds.

The paired CPTAC data also allowed RNA and protein associations with cell-state programs to be compared in matched patients. EGFR was positively associated with bulk pEMT at both the RNA and protein levels. The pEMT association was more negative for ERBB3 RNA than for HER3 protein in the same patients, while TACSTD2 RNA and TROP2 protein showed no clear bulk pEMT association (Fig. 6G).

Differentiation-1 associations were evaluated in the same within-target patient sets as pEMT. ERBB3 RNA and HER3 protein were both positively associated with differentiation-1 in the 84 paired patients (Spearman ρ, 0.470 and 0.345, respectively), as were TACSTD2 RNA and TROP2 protein in 108 patients (ρ, 0.598 and 0.467; Fig. 6H). The ERBB3/HER3 RNA-minus-protein correlation difference was 0.125 (95% CI, −0.118 to 0.371). These between-tumor bulk associations extend the differentiation-related target profiles to both RNA and protein measurements.

An external GeoMx analysis included 60 patients with one tumor region per patient matched across serial-section RNA and antibody-signal assays (GSE290057/GSE288406). Of 15 RNA-upper-quartile cases per target, eight were also in the EGFR antibody-signal upper quartile, five in the HER3 upper quartile and eight in the TROP2 upper quartile (Supplementary Fig. S12A–F). These are assay-signal comparisons: HER3 and TROP2 signals did not exceed the maximum of five IgG controls in 39 and 29 of 60 regions, respectively, and this background criterion was not a validated LOQ.

Available-case CPTAC analyses of both differentiation programs retained 108 RNA measurements and 84 HER3 protein measurements (Supplementary Fig. S13A,B). In 28 CODEX laryngeal SCC cores, the EGFR association with a LAMC2/podoplanin proxy was positive using intensities (partial ρ, 0.245; 95% CI, 0.180–0.311) but negative using Nimbus scores (−0.098; 95% CI, −0.155 to −0.040); several control intensities were also positively associated with the intensity-based proxy. In eight protein-complete HNSCC cell lines, the protein-balance index correlated with RNA-derived pEMT at 0.57 (95% CI, −0.26 to 1.00; q = 0.30), and at 0.41 in the lineage-adjusted analysis of 28 protein-complete squamous lines (95% CI, 0.06–0.67; q = 0.14; Supplementary Fig. S14). The cell-line estimates did not meet the multiplicity-adjusted significance threshold. Overall, paired tumor data linked EGFR to pEMT and HER3/TROP2 to differentiation at both RNA and protein levels.

### Bulk tumors recapitulate the pEMT-associated target-transcript balance

To test whether the relative transcript pattern extended to a larger tumor population, we analyzed bulk TCGA-HNSC tumors. Of 520 primary tumors, 277 met the specified HPV-assay eligibility rule and 273 had ABSOLUTE purity estimates available for molecular analysis (Fig. 7A; Supplementary Fig. S15A,B). Eligibility required at least one negative result and no positive result among p16 immunohistochemistry, HPV in situ hybridization and the TCGA RNA-seq HPV call; 21 eligible cases were HPV-DNA-positive by MassArray, which was not an eligibility assay. Target-transcript balance was summarized as I = z(EGFR) − ½[z(ERBB3) + z(TACSTD2)].

**Figure 7.**
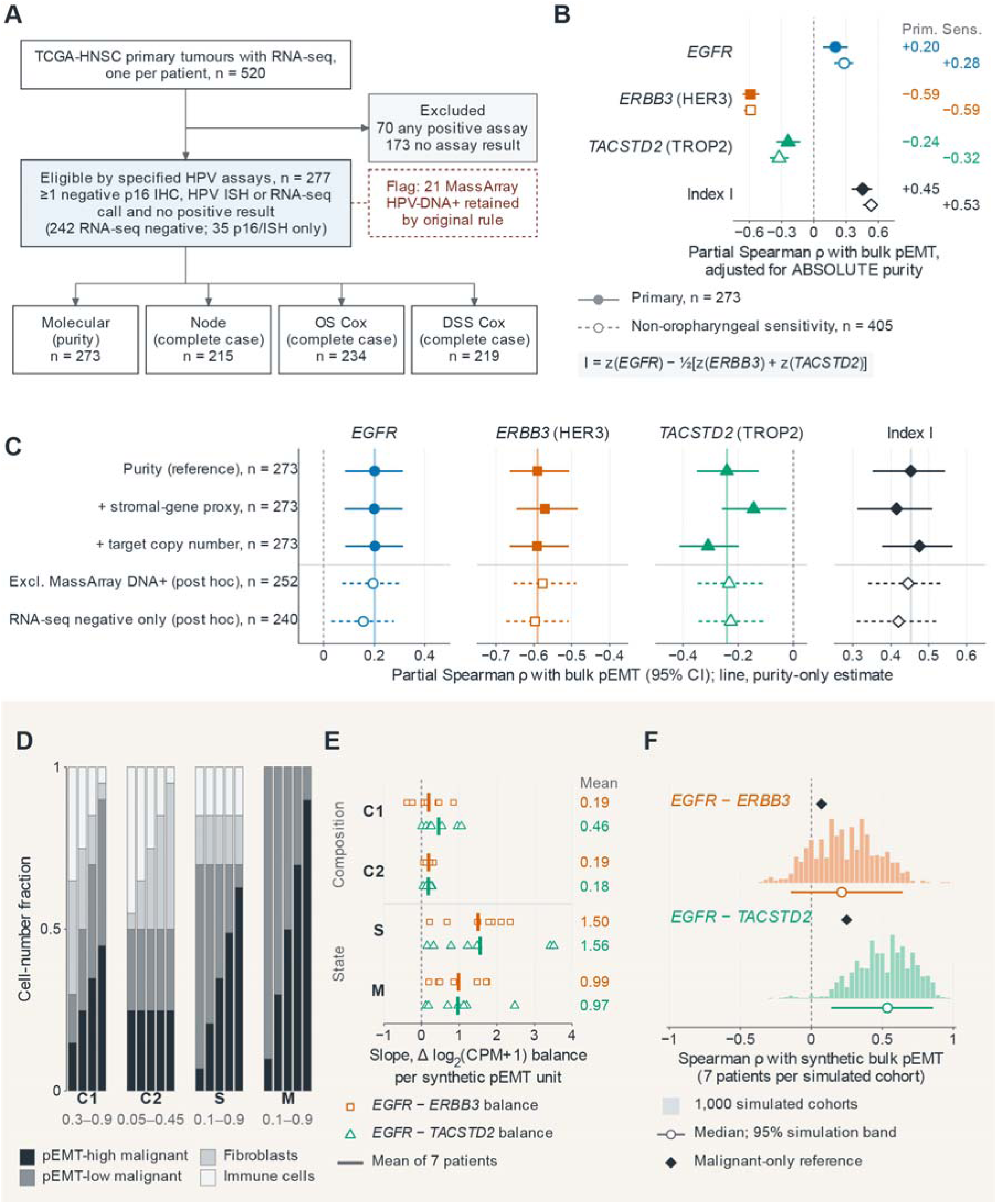
Bulk target-transcript balance tracks pEMT in TCGA-HNSC, with synthetic state and composition simulations. (A) Flowchart of TCGA-HNSC cohort selection and analysis populations. (B) Partial Spearman ρ of individual targets and a combined Index *I with bulk pEMT, adjusted for tumor purity. Filled and open symbols indicate primary and sensitivity cohorts. (C) Primary correlations with additional adjustments (stromal proxy, copy number) or restricted subsets. (D) Cell-number fractions of four synthetic pseudobulk designs (C1, C2, S, M). (E) Within-patient slopes of target balances (EGFR−ERBB3 and EGFR−TACSTD2) across synthetic pEMT units. (F) Distributions of Spearman* ρ *for target balances with synthetic bulk pEMT across 1,000 simulated cohorts*.

The balance index was positively associated with bulk pEMT after adjustment for purity (partial Spearman ρ, 0.453; 95% CI, 0.353–0.543; n = 273). Individual-target correlations were 0.201 for EGFR, −0.590 for ERBB3 and −0.241 for TACSTD2 (all q < 0.001; Fig. 7B). A broader non-oropharyngeal sensitivity population retained cases with unknown HPV status but excluded positive results on the specified HPV assays. It showed the same directions, with an index correlation of 0.53 among 405 tumors with purity estimates (Fig. 7A,B).

The index association was retained across the specified adjustment analyses and post hoc eligibility sensitivities. Additional adjustment for a stromal-gene expression proxy or target copy number yielded correlations of 0.415 and 0.475, respectively. Excluding MassArray HPV-DNA-positive cases or restricting the analysis to tumors with an RNA-seq HPV-negative call gave correlations of 0.446 and 0.420 (Fig. 7C).

Synthetic pseudobulks from seven discovery patients examined how malignant-cell state and cellular admixture could contribute to such patterns (Fig. 7D). At fixed cell-type proportions, varying the pEMT-high share of malignant cells produced positive mean slopes of 1.50 and 1.56 for the EGFR−ERBB3 and EGFR−TACSTD2 log (CPM + 1) balances, respectively; malignant-only mixtures gave slopes of 0.99 and 0.97. Varying the malignant-cell fraction at a fixed 1:1 pEMT-high-to-low ratio also produced positive slopes (0.19 and 0.46), as did varying the fibroblast fraction (0.19 and 0.18; Fig. 7E).

Across 1,000 composition-only simulated cohorts, the median between-patient correlations were 0.21 and 0.54 for the EGFR−ERBB3 and EGFR−TACSTD2 balances, respectively (Fig. 7F). These simulations show that both malignant-cell state and cellular admixture can produce the observed direction of target-balance variation; they do not quantify their respective contributions to the TCGA association.

The specified clinical models did not establish an association with nodal involvement or survival. Per standard-deviation increase in I, the adjusted odds ratio for pathological nodal involvement was 1.13 (95% CI, 0.85–1.52; n = 215), and the hazard ratios for overall and disease-specific survival were 1.20 (95% CI, 0.99–1.46; n = 234) and 1.13 (95% CI, 0.87–1.47; n = 219), respectively (q = 0.40, 0.18 and 0.40; Supplementary Fig. S15C). Post hoc subsite-stratified results remained exploratory; clinical-model sensitivities, the descriptive survival display and proportional-hazards diagnostics are shown in Supplementary Fig. S15C–E. Together, TCGA extends the target-transcript pattern to a larger tumor population, while the simulations show why bulk associations require joint consideration of malignant-cell state and tissue composition.

## Discussion

We identify a reproducible difference in how EGFR, ERBB3 and TACSTD2 transcripts relate to malignant-cell states in HNSCC. Within individual tumors, higher pEMT was associated with a relative shift toward EGFR, whereas epithelial differentiation showed a more positive relationship with ERBB3 and TACSTD2 than with EGFR. The pEMT-associated contrasts replicated in a separate single-cell cohort and were directionally concordant in two additional cohorts. Spatial profiling extended the relative EGFR pattern to tumor regions and linked the pEMT-high, less differentiated phenotype to histologically defined tumor buds. Paired proteogenomic measurements provided complementary evidence, connecting EGFR to pEMT and HER3/TROP2 to epithelial differentiation at both RNA and protein levels across tumors. Together, these findings place cell state alongside overall abundance as an important dimension of target-expression heterogeneity.

The central contribution of this study is a direct comparison of candidate targets within the same tumor. Previous single-cell studies established pEMT and epithelial differentiation as major sources of malignant-cell heterogeneity in HNSCC, while quantitative tissue assays have demonstrated broad, variable expression of EGFR, HER3 and TROP2 across tumors^6,7^. Our analyses connect these observations by asking whether the three target transcripts vary together across malignant-cell states. Broad detection and state-associated variation proved compatible: TACSTD2 was the most widely detected target transcript in the discovery atlas, yet its relative expression profile differed consistently from that of EGFR across pEMT states (Figs. 1D and 2C). A target can therefore be broadly represented in a tumor while still varying meaningfully among its malignant-cell populations. This distinction adds information that a tissue-average expression value or a binary detection category does not capture.

The patient-paired design makes this comparison particularly informative. Each target was evaluated across the same within-patient state groups, and the differences between target-specific effects were tested directly. This addresses a comparative biological question rather than inferring divergence from separate significance tests. Excluding the target genes from the state programs avoided direct inclusion of the measured targets in their explanatory scores, while equal-patient summaries kept tumors with more recovered cells from dominating the results. Both EGFR-relative pEMT contrasts were positive in every evaluable discovery and replication patient; in the exploratory synthesis excluding discovery, the corresponding contrasts were positive in all 11 and all 15 evaluable patients (Fig. 3A). The consistency of the relative ordering across independently generated datasets is a major strength, particularly because the absolute expression profiles and target detectability varied between cohorts.

Considering epithelial differentiation alongside pEMT gives the target differences a more specific biological interpretation. The two programs were related but not interchangeable, and differentiation-1 showed a relative target pattern opposite to that observed across pEMT states (Fig. 3B,D). Adjustment for differentiation attenuated the EGFR–TACSTD2 contrast, particularly in the replication cohort, indicating that this separation shares substantial variation with epithelial differentiation. The EGFR–ERBB3 contrast retained a positive direction in every discovery and replication patient after adjustment (Fig. 3C). These results favor interpreting the three targets in relation to both programs rather than assigning each target to a single epithelial–mesenchymal axis. For TACSTD2, the differentiation-associated pattern is consistent with experimental work linking TROP2 loss to mesenchymal features in squamous carcinoma^11^. Our findings extend that context to variation among malignant cells within individual tumors and suggest that differentiation markers may be useful companions to TROP2 assessment.

The relative association of EGFR with pEMT also fits existing functional evidence. Puram et al. localized pEMT-high malignant cells to the leading edge of HNSCC, and Zhou et al. identified an EGFR-associated regulatory network involved in local invasion^7,10^. Our study adds a comparative dimension: within the same tumors, the pEMT association of EGFR exceeded that of both partner targets. HER3 requires a different distinction between expression and activity. Experimental HNSCC models have shown that TROP2 loss can promote neuregulin-dependent HER3 activation. Lower ERBB3 transcript abundance in a pEMT-high population is therefore compatible with biologically relevant HER3 signaling^12^. This provides a rationale for future studies to measure receptor abundance and activation separately when testing how these target profiles relate to epithelial plasticity.

The spatial analyses connect these transcriptional relationships to tissue organization. In pathologist-annotated SCC regions, the relative EGFR–pEMT association remained positive after adjustment for composition proxies, sequencing depth and differentiation, and was maintained across alternative analytical settings (Fig. 4D–G). The paired tumor-bud analysis addressed a complementary question by linking the state programs to a recognizable histological compartment. Building on the original spatial characterization of this cohort, our analysis placed the buds within a joint pEMT-high, low-differentiation context in most patients (Fig. 5A–C)^9^. Their target profiles were not a uniform increase or decrease: the mean bud-minus-bulk EGFR signal was close to zero, whereas the mean ERBB3 and TACSTD2 signals were lower (Fig. 5D). Because most bud ERBB3 measurements were below the quantification limit, that comparison is best interpreted as a regional signal difference. The clearest contribution of the budding analysis is its histological anchoring of the joint pEMT-high, low-differentiation phenotype in which target heterogeneity can be studied.

Paired RNA and protein measurements add an important molecular perspective. EGFR and TROP2 showed close RNA–protein correspondence across CPTAC tumors, with correlations of 0.862 and 0.764, respectively, compared with 0.404 for HER3 (Fig. 6A–E). Beyond these overall correlations, the matched analyses linked EGFR positively to bulk pEMT at both molecular levels and linked HER3 and TROP2 positively to differentiation-1 (Fig. 6G,H). Thus, the differentiation-related target profiles identified in the transcriptomic analyses also had a protein-level counterpart across tumors. The pattern was target-specific: TROP2 showed a clearer association with differentiation than with bulk pEMT, and the negative pEMT association of ERBB3 RNA was weaker for HER3 protein. This argues for retaining target-specific information when moving from transcriptional profiling to protein-based biomarker development. In particular, direct HER3 measurement would add information that an RNA-only assessment captures less closely, while the EGFR and TROP2 concordance supports using RNA profiling to guide subsequent protein-focused investigation.

The bulk analyses extend the relative transcript pattern to a larger tumor population and clarify how it should be interpreted. In TCGA-HNSC, the target-balance index remained associated with pEMT after adjustment for purity and in analyses incorporating stromal expression, target copy number and alternative HPV eligibility rules (Fig. 7B,C). The synthetic pseudobulks then examined two distinct sources of this association. Varying malignant-cell state at fixed cell-type proportions produced the expected shift in target balance, including in malignant-only mixtures; varying tissue composition could also produce same-direction associations (Fig. 7D–F). These simulations offer a constructive link between the single-cell and bulk results by showing how both state and composition can influence a tissue-level measurement. This is consistent with previous work separating malignant-cell EMT from stromal expression programs. The combined evidence supports interpreting bulk target profiles with information on both cellular composition and differentiation, rather than treating a whole-tumor pEMT score as a direct measure of malignant-cell state^14^.

These findings provide a biological basis for incorporating cell-state measurements into translational studies of EGFR-, HER3- and TROP2-directed therapies. Clinical activity has been reported for TROP2-directed sacituzumab govitecan and EGFR-directed becotatug vedotin in previously treated HNSCC, and EGFR–HER3 bispecific ADCs have entered clinical evaluation in advanced solid tumors. Our results motivate asking where the relevant target proteins are expressed within a tumor, in addition to how much is present overall^3–5^. A focused next step would combine quantitative membrane EGFR, HER3 and TROP2 measurements with independent pEMT and differentiation markers in matched tumor regions, including buds and larger tumor nests. Such measurements could test whether the transcript-level contrasts correspond to differences in accessible antigen across cell states. State-characterized models could then evaluate internalization and drug response, including whether multi-specific targeting addresses populations with different target profiles. This sequence would connect the present biological observations to treatment-specific biomarker questions without relying on a universal expression threshold^46^.

The main boundaries of the study concern the scope of inference and the next stage of translation. The analyses are cross-sectional reanalyses of public cohorts, with modest patient numbers in individual single-cell datasets and differences in sampling, HPV ascertainment and assay platforms; several extensions were exploratory. Spatial measurements resolve multicellular regions, and CPTAC quantifies bulk protein rather than membrane abundance in individual malignant cells. The target-balance index was not established as a predictor of nodal involvement or survival, and the analyzed measurements were not linked to ADC response. These considerations define the priorities for validation: larger prospectively collected cohorts, spatially resolved surface-protein measurements and treatment-linked sampling. The replicated within-patient contrasts provide a specific set of hypotheses for those studies.

Overall, this study identifies cell-state-associated differences among three therapeutically relevant target transcripts in HNSCC and places them in spatial, histological and proteogenomic context. Its principal advance is the reproducible relative association of EGFR with pEMT, considered alongside the differentiation-associated profiles of ERBB3/HER3 and TACSTD2/TROP2. This integrated view provides a foundation for evaluating therapeutic targets in the malignant-cell populations and tissue compartments in which they are expressed, potentially guiding the rational design of target-specific or bispecific antibody-drug conjugates.

## Conclusion

This study reveals a reproducible divergence in target–state associations in HNSCC: within individual tumors, pEMT-high states showed a relative shift in transcript balance toward EGFR compared with ERBB3 and TACSTD2, whereas epithelial differentiation showed the opposite relative pattern. Spatial profiling supported the regional EGFR–pEMT association, tumor-bud analyses linked pEMT-high states to reduced differentiation, and paired bulk RNA–protein analyses provided complementary evidence for target-specific state associations. Together, these findings highlight malignant-cell state as an important dimension of therapeutic-target heterogeneity beyond overall abundance. They provide a biological rationale for integrating target-protein measurements with independent cell-state markers in prospective biomarker studies within clinical trials of EGFR-, HER3- and TROP2-directed antibody–drug conjugates.

## Declarations

### Ethics approval and consent to participate

Not applicable. This study exclusively reanalyzed publicly available, de-identified human datasets generated by previously published studies and public consortia. Ethical approval and informed consent were obtained by the original investigators in each respective study.

### Availability of data and materials

All human datasets are publicly available and de-identified. Data were obtained from the following repositories: scRNA-seq: GEO (GSE181919, GSE172577, GSE188737, GSE182227). Spatial transcriptomics: GEO (GSE208253, GSE300414); matched GeoMx RNA/protein data: GEO (GSE290057, GSE288406). CODEX imaging: BioImage Archive (S-BIAD3612). CPTAC-HNSCC: CPTAC portal. CCLE/DepMap: DepMap portal. TCGA-HNSC: UCSC Xena and PanCanAtlas. The reproducible code archive is available at Zenodo (https://doi.org/10.5281/zenodo.23655066).

### Authors’ contributions

Y.C. and P.Y. contributed equally to this work. Y.C. developed the statistical and computational methodology, curated the public datasets, performed the single-cell, spatial, proteogenomic, bulk transcriptomic, and synthetic pseudobulk analyses, generated the figures, maintained the reproducible code archive, and wrote the original draft of the manuscript. P.Y. contributed to data analysis, validation, interpretation of the results, and drafting and revision of the manuscript. J.L. and H.Y. contributed to data analysis and provided analytical advice and critical comments on the study. N.J. conceived and led the project, supervised the study, provided critical scientific feedback and biological interpretation, and critically revised the manuscript for important intellectual content. L.C., W.T., and J.J.T. supervised the study, provided critical scientific feedback, and critically revised the manuscript for important intellectual content. All authors discussed the results, read, and approved the final manuscript.

## Acknowledgements

We thank the investigators and participants of the original studies.

## List of abbreviations

ADC: Antibody-drug conjugate
AOI: Area of illumination
BH: Benjamini-Hochberg
CCLE: Cancer Cell Line Encyclopedia
CI: Confidence interval
CNV: Copy-number variation
CODEX: Co-detection by indexing
CP10K: Counts per 10,000 unique molecular identifiers
CPM: Counts per million
CPTAC: Clinical Proteomic Tumor Analysis Consortium
DAPI: 4’,6-diamidino-2-phenylindole
DSS: Disease-specific survival
EGFR: Epidermal growth factor receptor
EMT: Epithelial-mesenchymal transition
GEO: Gene Expression Omnibus
H&E: Hematoxylin and eosin
HER3: Human epidermal growth factor receptor 3
HNSCC: Head and neck squamous cell carcinoma
HPV: Human papillomavirus
HR: Hazard ratio
IgG: Immunoglobulin G
IHC: Immunohistochemistry
ISH: In situ hybridization
LOQ: Limit of quantification
MS1: Precursor-ion mass spectrometry
NB2: Quadratic negative binomial
OR: Odds ratio
OS: Overall survival
pEMT: Partial epithelial-mesenchymal transition
Q3: Upper-quartile (75th-percentile) normalization
QC: Quality control
RSEM: RNA-Seq by Expectation-Maximization
SCC: Squamous cell carcinoma
TCGA: The Cancer Genome Atlas
TPM: Transcripts per million
TROP2: Trophoblast cell-surface antigen 2
UMAP: Uniform manifold approximation and projection
UMI: Unique molecular identifier
WES: Whole-exome sequencing

